# Differential integration of activation and repression signals in a multi-enhancer system

**DOI:** 10.1101/2022.05.12.491703

**Authors:** Peter H. Whitney, Bikhyat Shrestha, Jiahan Xiong, Tom Zhang, Christine A. Rushlow

**Affiliations:** Department of Biology, New York University, New York, NY 10003, USA

**Author notes:** These authors contributed equally (Bikhyat Shrestha, Jiahan Xiong).

**Keywords:** shadow enhancers, MS2 live imaging, transcriptional kinetics, morphogen gradient

## Abstract

Transcription in the early *Drosophila* blastoderm is coordinated by the collective action of hundreds of enhancers. Many genes are controlled by so-called “shadow enhancers,” which provide resilience to environment or genetic insult, allowing the embryo to robustly generate a precise transcriptional pattern. Emerging evidence suggests that many shadow enhancer pairs do not drive identical expression patterns, however the biological significance of this remains unclear. In this study we characterize the shadow enhancer pair controlling the gene *short gastrulation* (*sog*). We removed either the intronic proximal enhancer or the upstream distal enhancer, and monitored *sog* transcriptional kinetics. Notably, each enhancer differs in *sog* spatial expression, timing of activation, and RNA Polymerase II loading rates. Additionally, modeling of individual enhancer activities demonstrates that these enhancers integrate activation and repression signals differently. While activation is due to the sum of the two enhancer activities, repression appears to depend on synergistic effects between enhancers. Finally, we examined the downstream signaling consequences resulting from the loss of either enhancer, and found changes in tissue patterning that are well explained by the differences in transcriptional kinetics measured.

**SUMMARY STATEMENT:** Non-intuitive shadow enhancer synergies are revealed by measuring transcriptional kinetics at the endogenous *short gastrulation* locus, giving rise to distinct patterning consequences in the dorsal ectoderm of *Drosophila* embryos.

## INTRODUCTION

*Drosophila* blastoderm development occurs rapidly over the course of 3 hours. During this time, all of the major tissue types are specified through a burst of intricate transcriptional regulation that culminates in the dramatic morphogenic events of gastrulation (reviewed in Stathopoulos and Newcomb, 2020). This period of development is a powerful system to study transcriptional regulation of developmentally relevant genes. In this study we explore the conserved phenomenon of “shadow” enhancers, first described in *Drosophila* (Hong et al., 2008). Enhancers are cis-regulatory elements that interact with transcription factors, and are capable of producing precise transcriptional outputs by employing a combinatorial logic of bound activators and repressors. Shadow enhancers have overlapping activities, that is, they activate transcription of the same gene in nearly identical patterns and are thought to provide robustness to the system (Frankel et al., 2010; Perry et al., 2010; Perry et al., 2011).

Further experiments have shown that shadow enhancers are widespread in developmentally relevant genes, and appear in multiple organisms, including humans (reviewed in Kvon et al., 2021). Gene editing and transgenic constructs have demonstrated that despite overlapping activities, RNA production from shadow enhancer pairs can deviate significantly, and multiple modes of enhancer interactions between shadow enhancer pairs have been identified. Shadow enhancers are said to have an *additive* interaction if the sum of the RNA produced from each individual enhancer matches what is produced from the wildtype enhancer pair. *Sub-additive* interactions are described as the sum of RNA produced being more than the wildtype RNA, while *super-additive* interactions describe the opposite. Finally, *repressive* interactions are the result of RNA from one enhancer exceeding the amount from the wildtype pair, suggesting that one of the enhancers is capable of repressing the output of the other (Kvon et al., 2021).

One of the first described shadow enhancer pairs was discovered at the *short gastrulation* (*sog*) locus (Hong et al., 2008). When cloned into transgenic expression constructs, both enhancers produce the characteristic lateral stripe of *sog* expression (Hong et al., 2008; Liberman and Stathopoulos, 2009). A recent report suggested that the one of the two enhancers may have repressive activity (Dunipace et al., 2019), and while particularly interesting at a mechanistic level, it is currently difficult to postulate a biological mechanism for how two enhancers both capable of driving expression can inhibit the total output of RNA. Therefore, we were motivated to quantifiably dissect exactly which features of transcription each enhancer controls, as solely measuring the total RNA produced obscures the multiple mechanistic steps involved in transcription. This would give a better understanding of transcriptional control by shadow enhancers more broadly, as the majority of genes active during early development have been shown to possess shadow enhancers.

To accomplish this, we first created several *Drosophila* lines with endogenous enhancer deletions to study the developmental consequences of abnormal *sog* expression. We inserted MS2 tags (see Methods) into the first intron of *sog* in all lines allowing us to compare transcription directly to wildtype alleles in fixed embryos, and to measure transcription in real time to examine how each enhancer modifies the parameters that define transcriptional output. We found that the *sog* enhancers have distinct but overlapping domains of expression, with individual enhancers capable of modifying different kinetic variables of transcription that combine in a manner that leverages the strength of each individual enhancer. This analysis also revealed that repression, but not activation, appears to be synergistic between the enhancers. Finally, we examined how altered transcription from the loss of individual enhancers leads to idiosyncratic downstream phenotypic consequences that are well explained by differences in the expression profile each enhancer alone generates.

## RESULTS

### Proximal and Distal enhancers of *sog* are together necessary to drive early blastoderm expression pattern

Early *sog* expression is controlled by two enhancers, originally known as “primary” or “intronic” and “shadow”, but also, and herein, referred to as “proximal” and “distal,” respectively (Dunipace et al., 2019; Hong et al., 2008). Fig. 1A shows the location of these two enhancers with respect to the transcription start site (arrow), with the distal enhancer located 20kb upstream (blue rectangle), and the proximal enhancer located approximately 1.5kb downstream within the first intron (green rectangle). Fig. 1B shows the location of key transcription factor binding sites in each enhancer that are largely responsible for the transcriptional domain of *sog*. Dorsal (Dl) serves as the primary transcriptional activator across the dorsal/ventral (D/V) axis while Zelda (Zld) potentiates Dl activity down the morphogen gradient, and Snail (Sna) represses activity in the mesoderm, resulting in the broad lateral stripes of the *sog* pattern (Liberman and Stathopoulos, 2009; Foo et al., 2014).

**Fig. 1.**
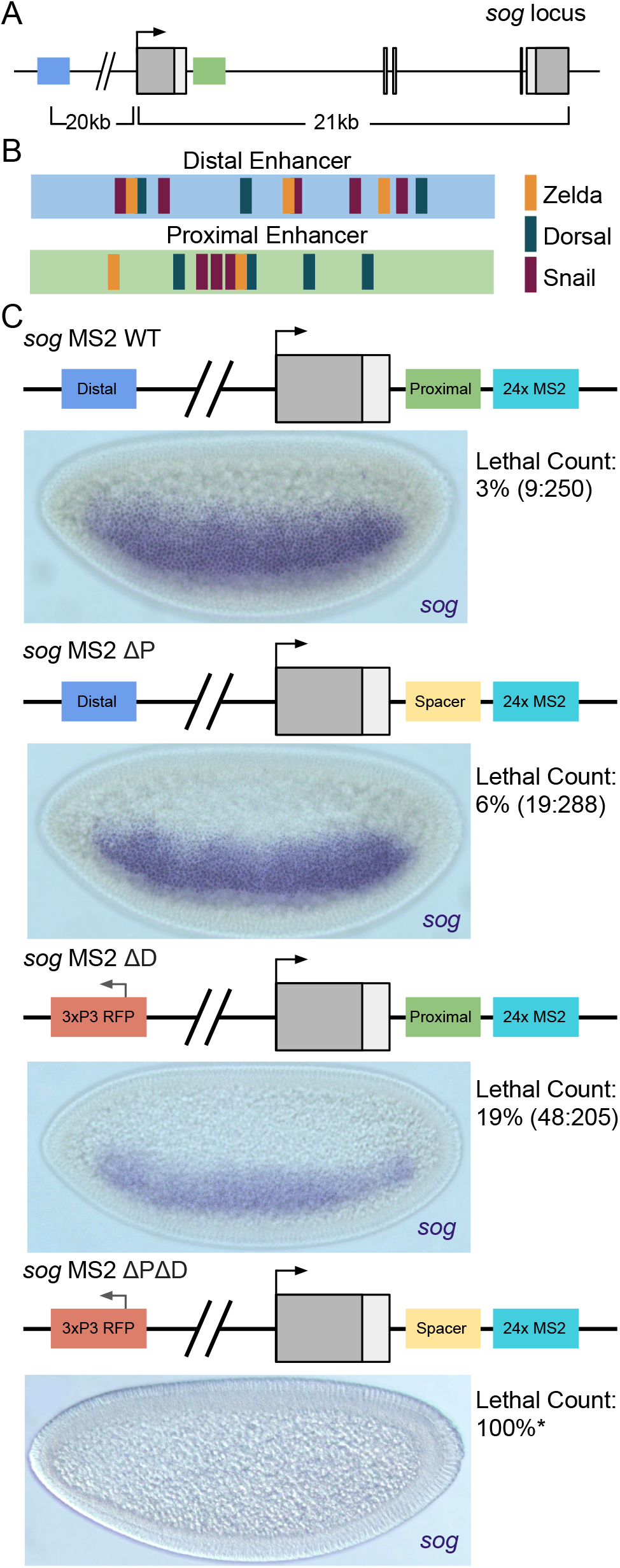
Early activation of *sog* is driven by two shadow enhancers. (A) Schematic representation of the *sog* locus. Previous studies have identified two enhancers that drive *sog* transcription (Dunipace et al., 2019). The proximal (green) enhancer located in the first intron of sog ∼2kb downstream of the promoter, and the distal enhancer (blue) located 20kb upstream of the promoter. (B) Transcription factor binding sites relevant to the expression of *sog*. Both enhancers contain binding sites for Zld (gold), Dl (dark green), and Sna (plum). All sites are present in roughly equal number, but vary in their position within each enhancer. (C) All enhancer lines created for this study. Each line contains a 1.2kb insertion of 24x MS2 loops located immediately downstream of the proximal enhancer. Δ*PsogMS2* and Δ*P*Δ*DsogMS2* replace the proximal enhancer with spacer DNA computationally depleted for early blastoderm transcription factor binding sites (Scholes et al., 2019) to maintain the spacing between the promoter and the MS2 loops. Δ*DsogMS2* and Δ*P*Δ*DsogMS2* replace the distal enhancer with a 3xP3 reporter construct for the purpose of screening mutant alleles. For each line, representative colorimetric *in situ* stainings for *sog* transcripts are shown in ventral lateral views. Lethal counts performed on all lines are listed besides each image. Δ*P*Δ*DsogMS2* produced no viable homozygous females or hemizygous males, and are therefore assumed to have a fully penetrant lethal phenotype.

To better understand the individual roles of these enhancers, we created enhancer deletion lines, in which we simultaneously inserted MS2 live-imaging tags within the first intron (Fig. 1C, turquoise rectangle) via CRISPR-Cas9 homology directed repair editing (see Methods). In order to maintain the spacing between the MS2 loops and the promoter in the proximal enhancer deletion, we adapted a “neutral” DNA sequence of identical size and GC content from Scholes et. al. (Scholes et al., 2019) to replace the proximal enhancer (Fig. 1C, yellow rectangles; see Methods). The wildtype enhancer allele, proximal enhancer deletion allele, and distal enhancer deletion allele will hereafter be referred to as *WTsogMS2*, Δ*PsogMS2*, Δ*DsogMS2*, respectively, and the double enhancer deletion allele as Δ*P*Δ*DsogMS2*.

To evaluate the fitness of our alleles, we performed lethal counts by counting the ratio of unhatched to hatched larvae from homozygous lines over a period of 36 hours (Fig. 1C, right). Both enhancer deletion lines showed increases in the number of unhatched larvae, with flies carrying Δ*DsogMS2* showing larger losses in viability than those with Δ*PsogMS2*. Δ*P*Δ*DsogMS2* failed to produce any homozygous flies, and therefore was assumed to be embryonic lethal. To evaluate the *sog* expression domains of these embryos, we performed colorimetric *in situ* hybridization for *sog* transcripts (see Methods). All alleles produced a *sog* expression pattern of varying intensity with the exception of Δ*P*Δ*DsogMS2*, which gave no apparent *sog* expression. We therefore concluded that both enhancers are necessary for *sog* expression, but a single enhancer is at least sufficient to generate some *sog* expression. In addition, *sog* does not appear to contain any other enhancers that drive early expression.

### Enhancer deletions appear to integrate position and output information separately

Because enhancers regulate gene expression at the level of transcription, we wanted to assess how each of the enhancers contribute to *sog* transcriptional output. To do this in a quantitative manner, we first turned to single molecule fluorescence in situ hybridization (smFISH), which is capable of producing fluorescence that scales linearly with the amount of RNA stained. We focused on measuring nascent transcripts, which can be seen in nuclei as large foci. To internally control for the dynamic nature of *sog* expression, particularly when ventral repression sets in during NC14, we crossed our MS2-tagged flies to wildtype flies to create *sog* heterozygous embryos. This allows us to directly compare both the level and domain of transcription of wildtype *sog* to enhancer deletion *sog*.

As diagrammed in Fig. 2A, the alleles can be discriminated through the use of two probe sets, a *sog* 5’ exonic-directed probe that labels both alleles in heterozygous flies (magenta), and a second probe set targeting the MS2 region and thus only the MS2 allele (cyan). This allows us to quantify transcriptional differences at the level of single nuclei (see Fig. 2B schematic of labeled foci). Fig. 2C shows the *sog* domain of a heterozygous *WTsogMS2*/wildtype embryo using this double labeling system in combination with an anti-Dl antibody. Note the *WTsogMS2* foci are white because of the dual labeling of magenta and cyan probes, while the wildtype foci are magenta since they are not labeled with the MS2 probe. For a more detailed explanation of allele discrimination and image analysis, see Fig. S1 and Methods.

**Fig. 2.**
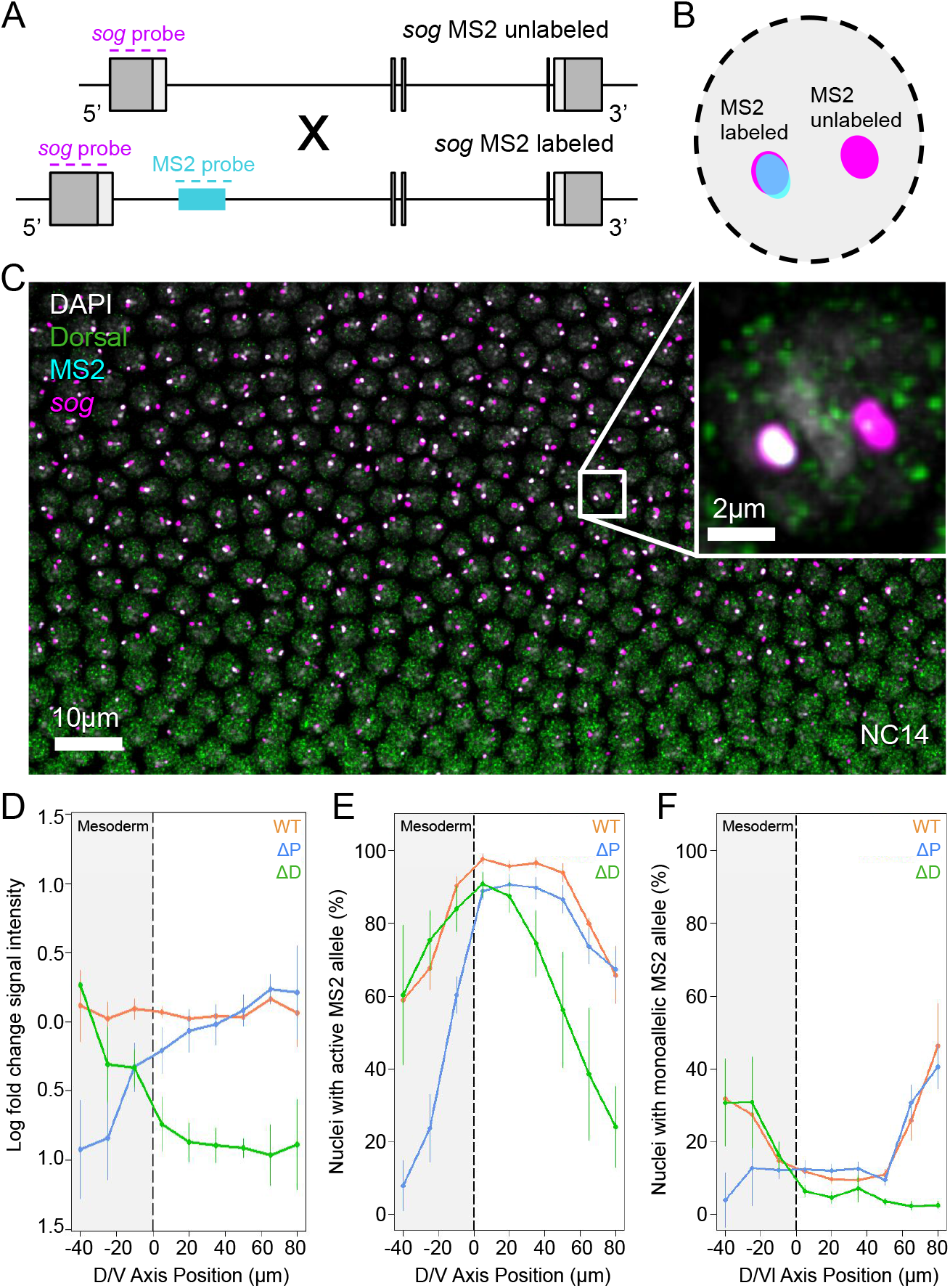
Internally controlled smFISH assay identifies spatial preference of each enhancer. (A) Crossing scheme used for all MS2 labeled lines. The location of exonic smFISH probe set (magenta) targets the first exon of *sog*, labeling both alleles, while the intronic smFISH probe set (cyan) targets only the MS2 sequence found in our engineered lines. (B) Schematic view of a single nucleus diagramming the expected allele labeling using the two probe sets. (C) Maximum intensity projection of z-stack images showing the region of the Dl gradient imaged. DAPI (white) labels nuclei, anti-Dl antibody (green) shows the Dl morphogenic gradient, MS2 probe (cyan) shows our MS2 tagged allele, and *sog* probe (magenta) shows all active *sog* transcription. Cut-out shows a single nucleus, matching the expectation of labeling in (B). (D) Log fold change calculated in each nucleus by taking the log ratio of the wildtype allele *sog* nascent transcript staining intensity over the MS2 allele *sog* nascent transcript staining intensity. Measurements were performed across the Dl gradient for *WTsogMS2* (orange), Δ*PsogMS2* (blue), and Δ*DsogMS2* (green). Shaded region with dashed line shows the location of the presumptive mesoderm. Error bars: s.e.m. (E) Quantification of the percentage of all active MS2 alleles regardless of the state of the wildtype allele. (F) Quantification of the percentage of all active MS2 alleles in nuclei with no detectable wildtype allele transcription.

The double labeling assay allows us to internally control for fluorescence of nascent transcripts by examining nuclei that have both alleles active and taking a log ratio between the intensity of the two magenta foci to look for upregulation or downregulation of *sog*, which will give positive or negative values respectively. To ensure that our *WTsogMS2* allele operates identically to our unlabeled wildtype allele, we plotted this ratio across the D/V axis and found minimal fluctuations around 0, indicating that there is no change in *sog* output with the addition of our MS2 tag (Fig. 2D, orange line). In contrast, Δ*PsogMS2* showed significant downregulation in the mesoderm but trended towards wildtype levels in the dorsal portion of the pattern (Fig. 2D, blue line), and Δ*DsogMS2* again displayed the opposite trend (Fig. 2D, green line). This suggests that the two enhancers regulate transcriptional output differentially active across the *sog* domain.

To further investigate the idea of enhancers having a spatial preference, we assessed the percentage of MS2-expressing nuclei in bins across the *sog* domain, including in the count not only nuclei with both alleles active, but also those with only the MS2 allele active (monoallelic expression). The *WTsogMS2* allele showed robust activation across the *sog* domain, with major reductions in activity at the ventral and dorsal extremes of the domain (Fig. 2E, orange curve). In contrast, the Δ*PsogMS2* allele showed a significant decrease in activation on the ventral end of the pattern (Fig. 2E, blue curve), and the Δ*DsogMS2* allele showed the opposite trend with significant decreases in the dorsal end of the pattern (Fig. 2E, green curve). Additionally, neither of the deletion alleles showed as robust an activation as the *WTsogMS2* allele in the lateral portion of the domain.

These results suggest that the transcriptional domain displayed by *sog* is the result of the two separate enhancers summing their individual domains, in a manner that displays simple additivity at the borders of the *sog* domain, and sub-additivity towards the center. This sub-additivity likely arises from a complete saturation of activation in the center of the pattern, i.e., there are simply no more nuclei to activate, rather than any particular transcriptional mechanism that causes the enhancers to integrate their activation signals in a fundamentally different way across the D/V axis. This simple framework is potentially broadly applicable across multiple shadow enhancer pairs, as it agrees with several previous studies where shadow enhancers appear to aid in creating robust borders to the transcriptional patterns they give rise to (Perry et al., 2011; El-Sherif and Levine, 2016; Dunipace et al., 2019; Scholes et al., 2019).

We then focused on only the monoallelic-expressing nuclei to determine if the occurrence of monoallelic expression was differentially influenced by either enhancer at any point in the *sog* domain. We found a bimodal distribution of MS2 monoallelic expression for the *WTsogMS2* allele, with peaks of monoallelic expression on both ends of the *sog* domain (Fig. 2F, orange curve). This is consistent with earlier reports that have suggested that monoallelic expression occurs more frequently on the border of transcriptional domains (Hoppe et al., 2020). When we examined the enhancer deletion lines, we found a similar trend to the previous experiment, where the dorsal peak of monoallelic expression vanished in our Δ*PsogMS2* (Fig. 2F, blue curve), and the ventral peak absent in the Δ*DsogMS2* allele (Fig. 2F, green curve). Strikingly, the peak that was not absent in each deletion allele remained at exactly the level we observed in wildtype (Fig. 2F, note region of overlap among the three curves). This suggests that monoallelic expression in shadow enhancer pairs may be largely driven by single enhancers acting alone. Taken together, these observations demonstrate that each of the *sog* enhancers has a preferred domain along the D/V axis, and when combined together create the wildtype pattern.

### MS2 live imaging shows shadow enhancers separately integrate kinetic properties of transcription

Next, we wanted to explore how control of transcriptional kinetics differed between the two enhancers. Measuring nascent transcription in fixed tissue confounds two critical variables, the timing of activation and the rate of transcript production. In order to examine these two variables, we turned to live imaging to visualize the number of nascent transcripts produced over time utilizing our endogenously inserted MS2 tag. Fig. 3A describes how MS2 live imaging operates with the intronically inserted MS2 loops, with MCP-GFP binding detectable only to transcripts that have not yet been spliced out of the first intron. Imaging was performed on the portion of the embryo that includes the *sog* expression domain from NC12 to mid-NC14, at which point the defined line of ventral repression in *sog* becomes apparent (see schematic in Fig. 3B, still images in Fig. 3C, and representative Movies S1-S3).

**Fig. 3.**
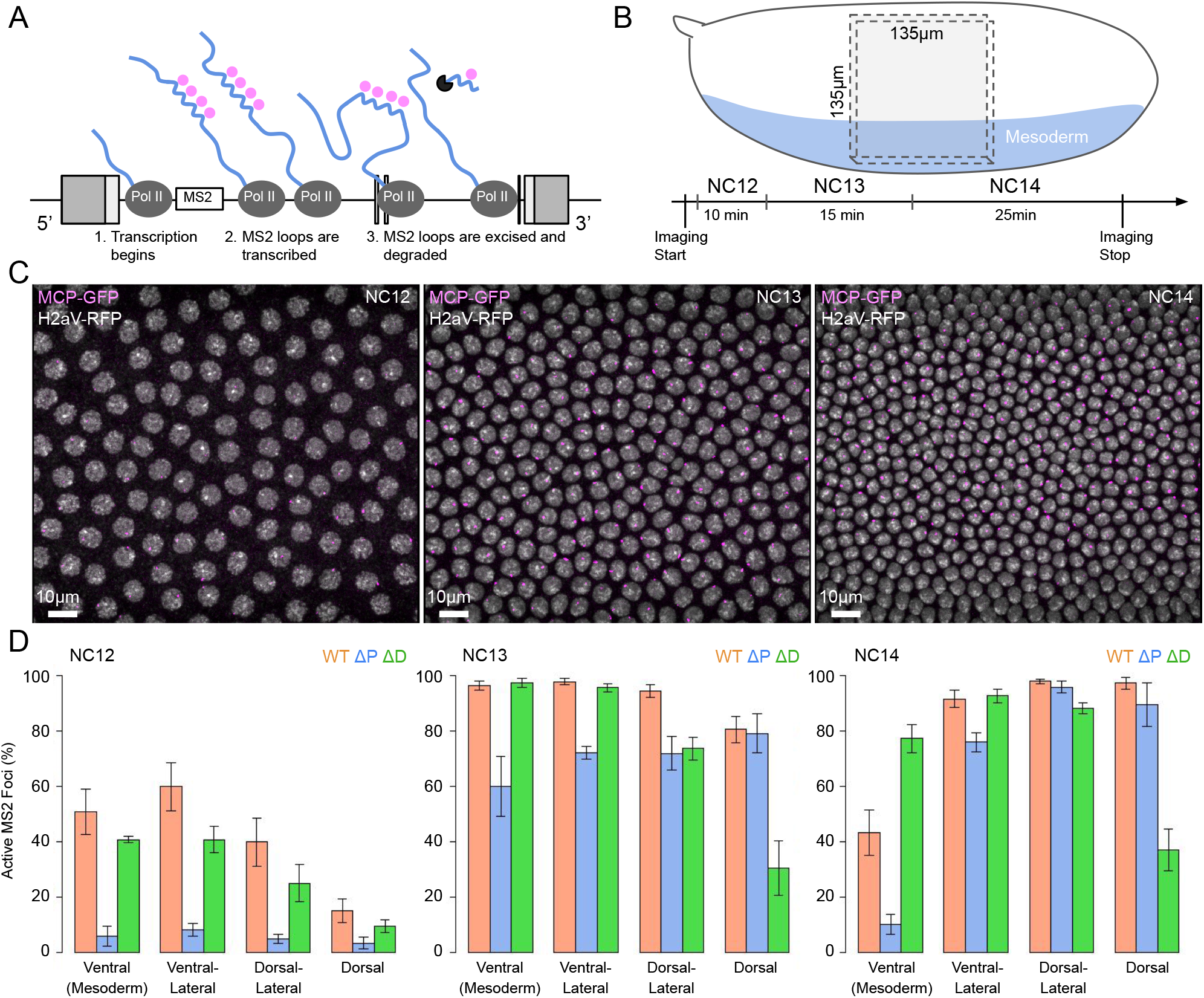
MS2 live imaging reveals differences in activation from NC12 to NC14. (A) Schematic of intronic MS2 loops reporting on live transcription. MS2 loops (blue hairpins) are transcribed and serve as binding sites for MCP-GFP (pink dots). Loops are spliced co-transcriptionally and are degraded by RNA-exonucleases (black circular sector). (B) Region of the embryo imaged during live imaging. Imaging volume of 135μm by 135μm by 15μm was positioned ventral/laterally to capture ventral repression as seen in late NC14 in order to orient nuclei across the D/V axis. Embryos were imaged for approximately 1 hour across NC12 to NC14. (C) Stills taken from live imaging movie of *WTsogMS2* Active transcription was determined by the appearance of MCP-GFP foci (pink) in nuclei marked by H2aV-RFP (white). Scale bar: 10μm. (D) Quantification of number of nuclei with active transcription for *WTsogMS2* (orange), Δ*PsogMS2* (blue), and Δ*DsogMS2* (green). Percentage of active nuclei were measured in the ventral region (mesoderm), ventral/lateral region, dorsal/lateral region, and dorsal region of the *sog* transcriptional domain. Error bars: s.e.m.

In order to characterize the *sog* transcriptional activity of each line and to validate the fidelity of our live imaging system, we counted the number of foci seen in each nuclear cycle relative to the number of nuclei (Fig. 3C). We classified foci position into four categories across the *sog* domain: ventral, ventral/lateral, dorsal/lateral, and dorsal, with the ventral position encompassing any foci detected in the presumptive mesoderm. Broadly, we observed that *WTsogMS2* and Δ*DsogMS2* produced similar numbers of MS2 active nuclei, with the exception of the dorsal most position (see Fig. 4D histograms). This is in contrast to the activity of Δ*PsogMS2*, which nearly universally underproduced relative to *WTsogMS2*. This is most striking in NC12, where barely any transcriptional activity was observed (Fig. 4D, blue bars). Furthermore, in NC14, Δ*PsogMS2* produced very little transcription in the ventral most bin, which suggests that the distal enhancer is more sensitive to Sna-mediated repression. The results of this analysis at NC14 are consistent with our fixed imaging data (Fig. 2D), demonstrating that the MS2 system is faithfully reporting on the transcriptional output of *sog*.

**Fig. 4.**
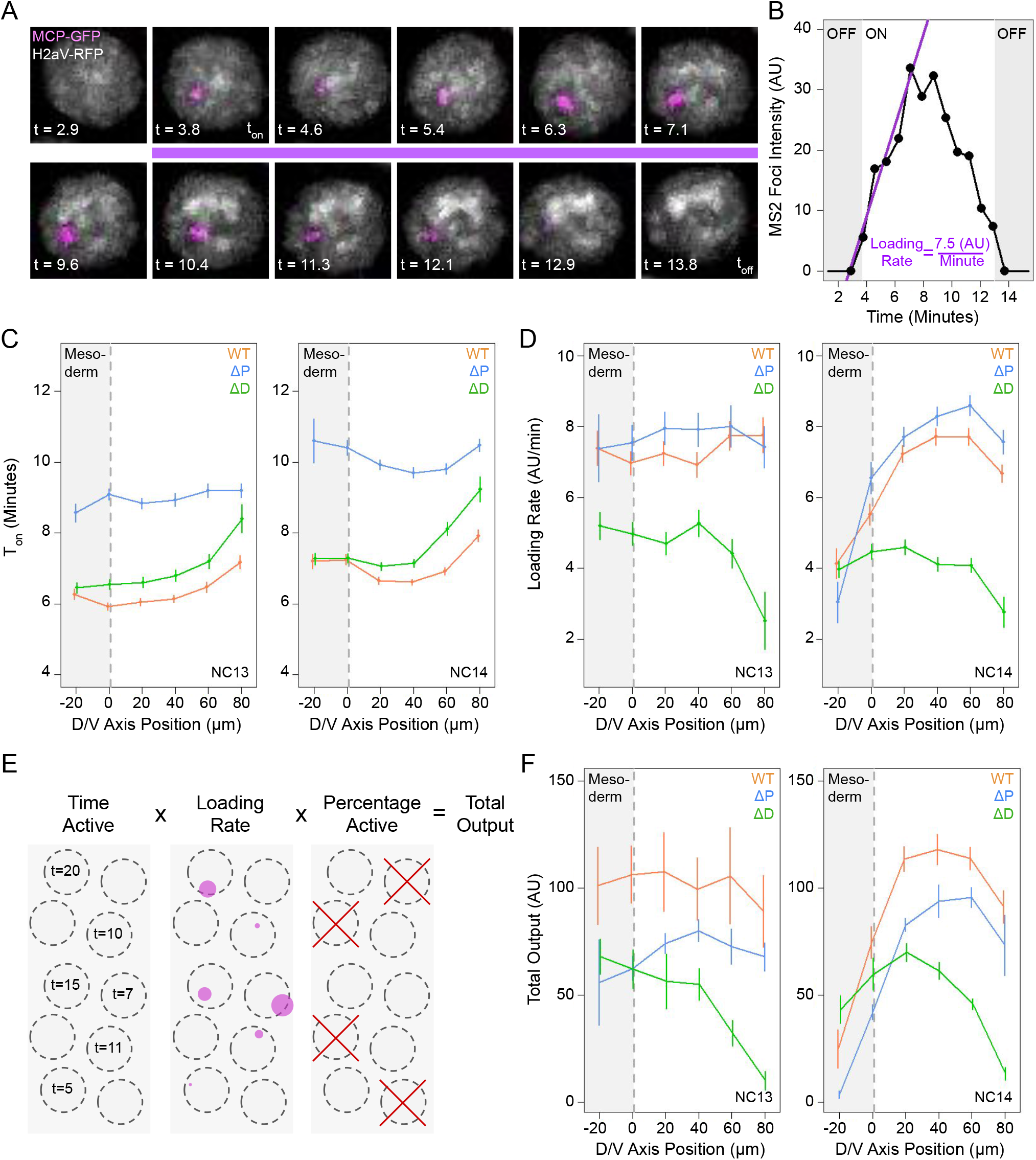
Internal kinetic parameters are modified by individual enhancers. (A) Maximum intensity projections of a single nucleus tracked over time. t_on_ is determined by the first appearance of a MCP-GFP focus (pinka) inside a H2aV-RFP labeled nucleus (white). The first 5 timepoints of a track focus (purple line) are used to determine the relative RNA Pol II loading rate. t_off_ represents the timepoint at which a focus can no longer be detected. (B) Signal intensity over time of the MCP-GFP focus tracked in (A). Loading rate is found by fitting a linear model (purple line) to the first five timepoints after t_on_. (C) t_on_ times for across the D/V axis for *WTsogMS2* (orange), Δ*PsogMS2* (blue), and Δ*DsogMS2* (green) at NC13 (left) and NC14 (right). Shaded region of the graph represents the mesoderm. Error bars: s.e.m. for all nuclei. (D) Relative loading rates measured across the D/V axis for all genotypes at NC13 (left) and NC14 (right). (E) Schematic diagram demonstrating how total transcriptional output is calculated. (F) Total output measured across the D/V axis for all genotypes at NC13 (left) and NC14 (right).

To better understand how activity differs between the enhancers, we analyzed single nuclei in the manner outlined in Fig. 4A-B. Single foci were tracked and their voxel intensity values summed for each timepoint to produce a trace of MS2 activity over time. Then, several parameters were extracted from these traces: t_on_, defined by the time at which a MS2 focus was first observed following the previous nuclear division; loading rate, which describes the rate of signal increase by fitting a line to values where the GFP signal first increases (Fig. 4B, purple line); and t_off_, the time at which the signal is no longer detectable in that nucleus. All parameters were measured for nuclei across the D/V axis. We focused on NC13 and NC14 for this analysis, as these cycles produce far more activity than NC12 and are therefore more relevant to the total transcriptional output of *sog*.

Fig. 4C shows the t_on_ times for each genotype at NC13 and NC14. With the exception of the most ventral bins, *WTsogMS2* activated transcription at a faster rate than both deletion genotypes. Δ*PsogMS2* showed extremely delayed transcription at all positions and times, in line with the results of inefficient activation discussed above. However, when we examined the loading rates of all lines (shown in Fig. 4D), Δ*PsogMS2* outperformed even *WTsogMS2* in most cases, and greatly outperformed Δ*DsogMS2*, which had loading rates that fell severely in the more dorsal bins. All genotypes showed lower loading rates in the ventral bins, likely driven by Sna-mediated repression of *sog*.

With these apparently opposing enhancer activities for activation and loading rates, we wanted to create a metric that would describe the total transcriptional output of each genotype. To do this, we adapted an approach used by Garcia et al. (2013) that described transcriptional output by combining multiple parameters of transcription (diagrammed in Fig. 4E). For each bin, we multiplied the time active (duration of t_on_; Fig. 4B, white area) by the loading rate for each nucleus. The average value obtained in each bin was then multiplied by the fraction of nuclei with detectable transcription to normalize the differences in activation seen across the genotypes. Total output values are plotted in Fig. 4F, showing that *WTsogMS2* generates the most activity by this metric. Thus, although at first glance it appeared that the distal enhancer when acting alone drives higher transcriptional activity than when the distal and proximal are combined (Fig. 4D; also shown by Dunipace et al. (2019), this is not the case when taking into account all transcription variables to determine total output.

Curiously, the only point *WTsogMS2* is not highest in this metric is in the most ventral bin in NC14, where Δ*DsogMS2* showed higher total output (see Fig. 4F, NC 14, compare orange and green lines in mesoderm). This result indicates a repressive interaction between the two enhancers, as an additive interaction is always indicated by the wildtype enhancer pair showing the highest output.

### Modeling the rate of activation predicts potential cross-talk of repression, but not activation

To address this finding, we wondered if it was possible to construct the observed wildtype transcriptional activation and repression of *sog* over time, using the kinetic parameters gathered from the enhancer deletion lines. By building a model of each individual enhancer’s activity over time, we could simulate what would be observed if those enhancers operated in the same nucleus, but did not interact when driving *sog* transcription. We could then compare the output of this simulation to the transcriptional activity observed in *WTsogMS2* embryos, where any significant deviations from the model’s prediction and the data could be interpreted as potential synergy between the enhancers.

In order to simulate enhancer activity, we first fit gamma distributions to the t_on_ and t_off_ values obtained for each genotype across the D/V axis at NC14. These distributions were then refined by systematically altering the shape and rate parameters of each distribution until the differences between simulated nuclei and the observed activity over time were minimized (see Methods). During simulation, distributions of t_on_ and t_off_ for each nucleus were sampled independently, assuming no correlation between a nucleus’ activation time and the time of loss of signal (see Fig. S2 for validation of this assumption).

Fig. 5A shows the output of simulations based on our model expressed as the percentage of active nuclei over time (solid line) plotted over the data gathered from our NC14 live-imaging experiments (open circles). Note the near perfect overlap for all genotypes, indicating the distributions of t_on_ and t_off_ values chosen are sufficient to describe the data. Beside each plot is shown the distributions of t_on_ (pink) and t_off_ (blue) values generated from sampling the fit gamma distributions. For a breakdown of distributions and fits for all D/V bins, see Fig. S3.

**Fig. 5.**
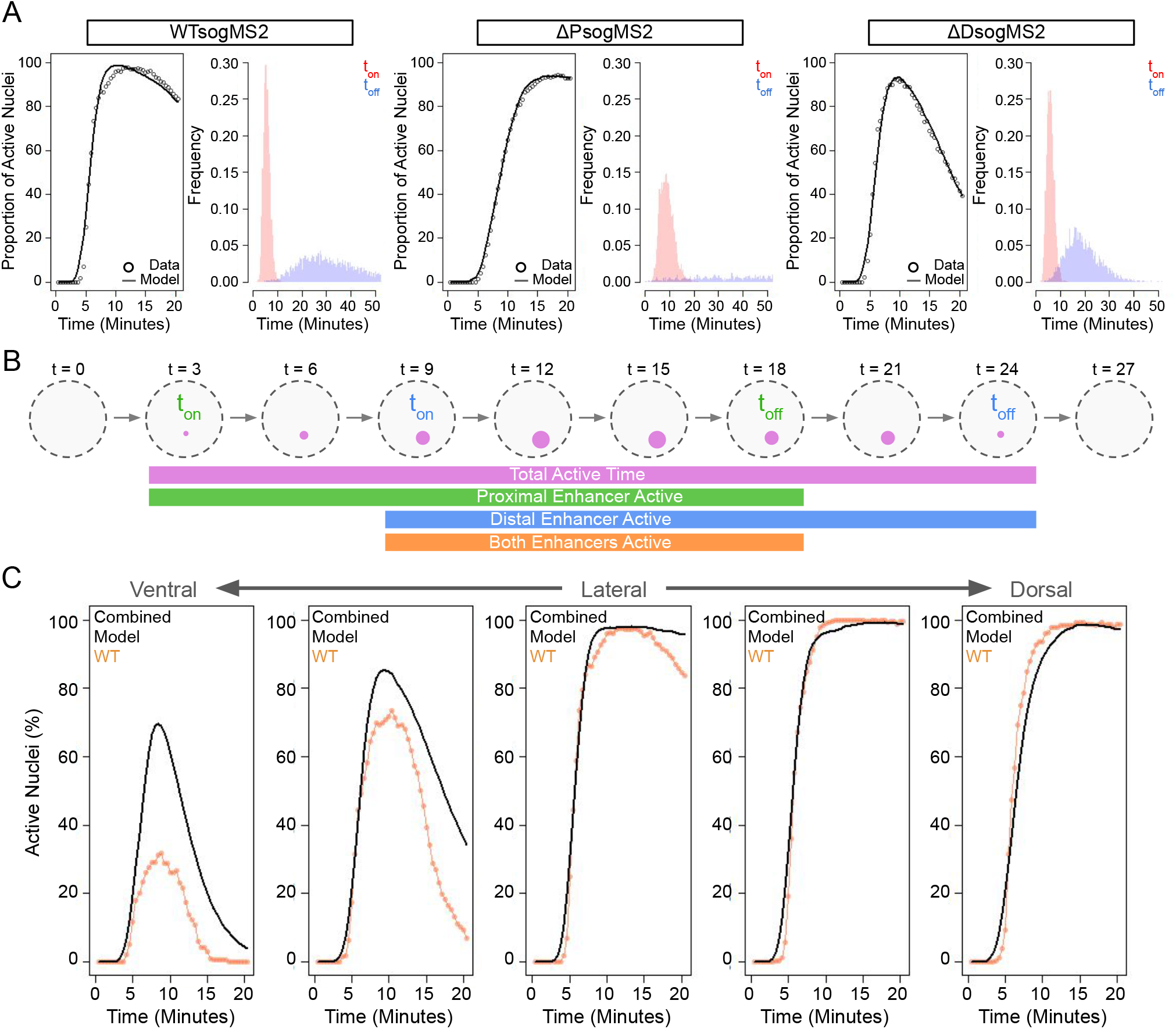
Modeling the activities of individual enhancers reveals potential synergy of Sna-mediated repression. (A) Activation over time of all genotypes in the lateral region of the embryo at NC14. Model fits (solid lines) based on simulations of 10,000 nuclei generated by sampling t_on_ and t_off_ distributions superimposed over data (open circles). Histograms of t_on_ (red) and t_off_ (blue) values used to perform simulations shown to the left. (B) Schematic representation of modeling *WTsogMS2* activation over time using t_on_ and t_off_ values from enhancer deletion distributions. Active transcription (purple foci) is maintained by the sequential and overlapping activity of individual enhancers. Enhancer activity (proximal in green, distal in blue) is defined by t_on_ and t_off_ values derived from each enhancer’s fit distributions. (C) Output of combined model of non-interacting enhancers (black line) compared to activation data from *WTsogMS2* (orange line). Each graph contains data from different spatial bins across the D/V axis.

Having found parameters for all distributions that can accurately describe the data of individual enhancers based on data from our enhancer deletion lines, we created a combined model that simulates the activity of both enhancers in a single nucleus. Nuclei remain “on” if at least one enhancer is simulated to be active based on the values obtained by sampling t_on_ and t_off_ distributions obtained from each deletion line. This underlying assumption represents the null hypothesis that there is no interaction between the enhancers, and the activity seen in *WTsogMS2* is based purely on the combined activity of the proximal and distal enhancers. Fig. 5B shows conceptually how the model interprets multiple sets of t_on_ and t_off_ values sampled from each pair of distributions for the two enhancer deletion genotypes. In this example, the faster acting proximal enhancer is responsible for the initial activation of *sog* (green), while the slower acting distal enhancer activates later (blue), with a brief period of overlapping activity of both enhancers (orange) that maintains continuity of transcription.

Using this combined model, we simulated an additional 10,000 nuclei for each bin across the D/V axis, and compared the results to the observed activation kinetics of *WTsogMS2*. While in all D/V bins the rate of activation was remarkably well predicted by the model (Fig. 5C, see overlap between initial rise in curves), the rate of deactivation was not, and a dramatic overactivation of the model output compared to the data was seen in the ventral bins (Fig. 5C, note different curve heights). This rigorously demonstrates that the strong repression experienced by the distal enhancer in the mesoderm is somehow influencing the ability of the proximal enhancer to activate transcription in *WTsogMS2* embryos. Additionally, it identifies the key parameter from which the repressive interaction arises, clearly implicating Sna-mediated repression, not Dl-activation. Understanding this form of crosstalk between enhancer pairs is likely critical for creating a unified model of enhancer biology. For a more detailed look at the implications of this finding and possible underlying mechanisms, see Discussion.

### *sog* Enhancer deletions affect downstream signaling events in late blastoderm embryos

With a better understanding of the kinetic features of *sog* transcription, we wanted to evaluate the downstream developmental effects that occur due to the loss of a single *sog* enhancer. To observe developmental consequences of the *sog* enhancer deletions, we measured the developmental morphogen gradient that Sog protein is directly involved in refining: the dorsally located gradient of phospho-Mothers Against Decapentaplegic (pMAD). As diagrammed in Fig. 6A, Sog protein produced in ventro-lateral cells diffuses dorsally, where it inhibits activity of the TGF-β homolog Decapentaplegic (Dpp), resulting in a gradient of Dpp (Shimmi et al., 2005; Wang and Ferguson, 2005). Dpp signal transduction leads to the phosphorylation of MAD, and in early NC14 it initially creates a broad region of pMAD. Sog protein binds to Dpp, preventing it from creating high levels of pMAD in the lateral regions of the embryo, and continued Sog diffusion eventually restricts pMAD to a Dl stripe 4-5 nuclei wide (Dorfman and Shilo, 2001; Rushlow et al., 2001; Sutherland et al., 2003). By staining embryos with anti-pMAD antibodies, we can visualize any impairments in pMAD-domain formation that may be caused by *sog* enhancer deletions.

**Fig. 6.**
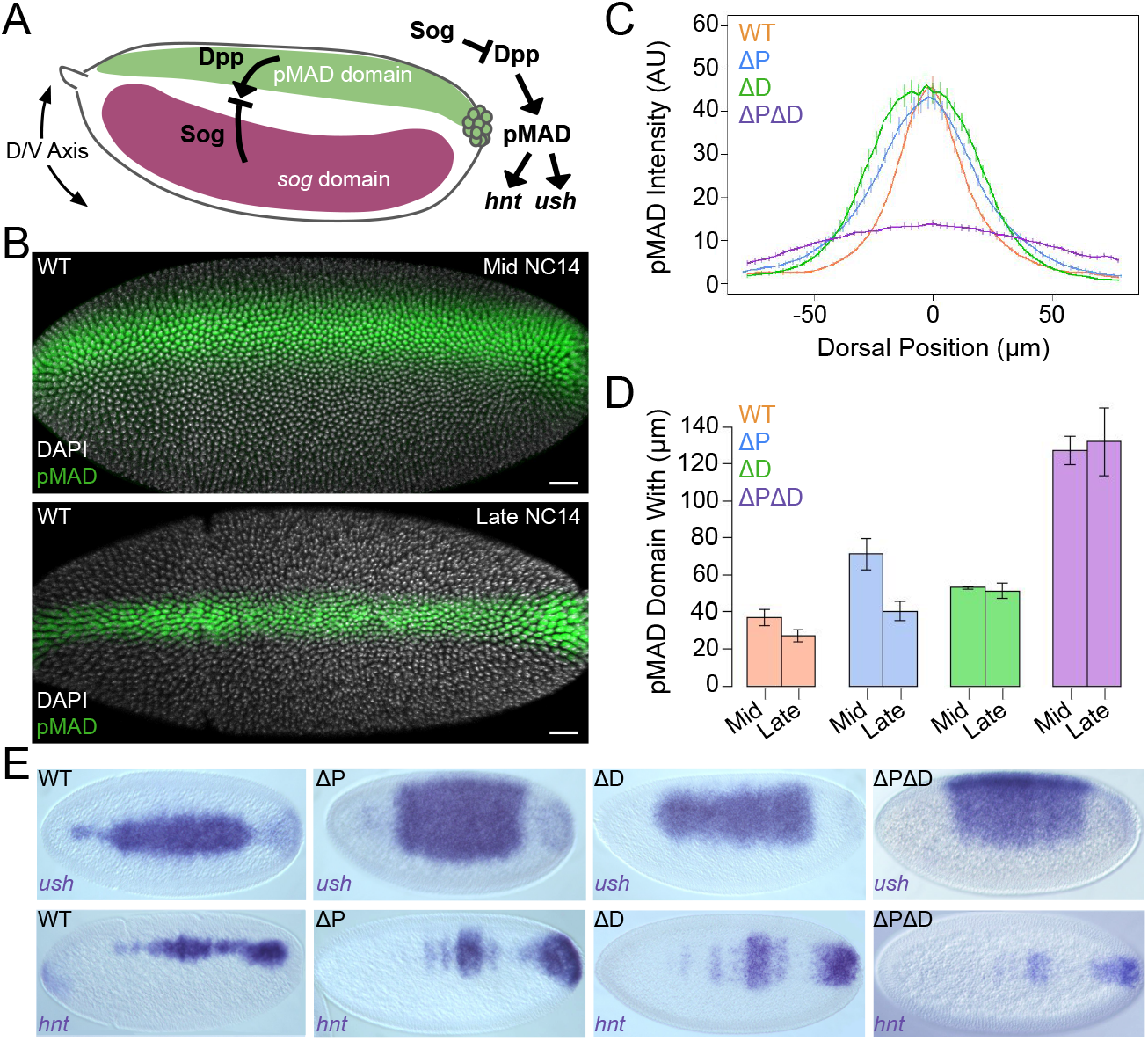
*sog* enhancer deletions show differential downstream effects on the pMAD gradient and pMAD target gene expression. (A) Schematic of the downstream signaling controlled by *sog*. Sog protein diffuses dorsally from the ventral-lateral *sog* domain (dark purple) where it encounters and sequesters ventrally diffusing Dpp emanating from the pMAD domain (green). Sog also localizes Dpp to the dorsal midline (Shimmi et al., 2005; Wang and Ferguson, 2005). Genetic interactions of the components of this pathway are shown to the right. pMAD acts as a transcription factor on target genes *hnt* and *ush*. (B) Dorsal views of mid and late NC14 homozygous *WTsogMS2* embryos stained with anti-1/5 pMAD antibody (green) and DAPI (white). Late embryos are identified by irregular nuclei shape and the appearance of the ventral furrow. Scale bars: 20μm. (C) pMAD staining intensity across the dorsal midline of the embryo for *WTsogMS2* (orange), Δ*PsogMS2* (blue), Δ*DsogMS2* (green) and Δ*P*Δ*DsogMS2* (purple). Each embryo is centered based on the point of highest pMAD intensity. Error bars: s.e.m. (D) Quantification of pMAD domain width for all genotypes in mid and late NC14 embryos. Domain width is determined by measuring the point at which pMAD staining intensity is above 50% of max intensity. Error bars: s.e.m. (E) Evaluation of pMAD target genes on all genetic backgrounds. Conventional colorimetric *in situ* hybridizations were performed on NC14 embryos. *ush* and *hnt* were chosen as representative early and late genes, respectively (Hoppe et al., 2020).

Fig. 6B shows the results of pMAD antibody staining on homozygous *WTsogMS2* mid and late NC14 embryos. Due to the continued production of Sog protein, we see a narrowing of the pMAD domain. To quantify total pMAD levels, we measured the intensity of pMAD staining and plotted it over the dorsal position centered on the peak of maximum pMAD staining (Fig. 6C). Interestingly, the Δ*DsogMS2* and Δ*PsogMS2* alleles show peak pMAD intensity nearly identical to wildtype, suggesting that only a small input of *sog* activity is required to increase the level of pMAD seen in the dorsal-most cells. This is in contrast to max pMAD levels seen in Δ*P*Δ*DsogMS2*, which completely fail to refine into a narrow peak. However, single enhancer deletions produced an overall broader distribution of pMAD staining. Δ*DsogMS2* embryos gave the broadest pMAD domain, which is consistent with the rank order of total output of *sog* (Fig. 4F).

Because we found large differences in the onset of transcription in our enhancer deletions, we were interested to see if this influenced the timing of pMAD refinement. To test this, we plotted the width of the pMAD domain of both mid and late NC14 embryos for all genotypes (Fig. 6D). As expected, *WTsogMS2* embryos refine their pMAD domain over these two timepoints. Δ*PsogMS2* embryos carry out a more extreme refinement, initially showing a far larger pMAD domain. In contrast, Δ*DsogMS2* embryos initially show a modestly expanded pMAD domain less so than Δ*PsogMS2* embryos, however this undergoes no appreciable change in late NC14. Finally, Δ*P*Δ*DsogMS2* displays an incredible expansion of the pMAD domain, which in the absence of any *sog* production, does not undergo any subsequent retraction.

These results are well explained by our MS2 data, which showed significant delays in the onset of transcription of *sog* in Δ*PsogMS2* embryos. However, the high loading rates achieved by the distal enhancer allow Δ*PsogMS2* embryos to eventually produce enough Sog protein to refine the pMAD gradient. The lack of refinement of pMAD in Δ*DsogMS2* is likely due to the inability of the primary enhancer alone to continuously produce *sog* transcripts late into NC14.

To determine if the changes in the pMAD gradient impact the expression domains of pMAD target genes, we performed colorimetric *in situ* hybridization for two representative pMAD targets; *u-shaped* (*ush*), thought to be an “early” pMAD target, and *hindsight* (*hnt*), thought to be a “late” pMAD target (Hoppe et al., 2020) (Fig. 6E). Patterns observed in *ush* stained embryos show Δ*PsogMS2* embryos more severely affected, creating broad domains of expression matching those found in the Δ*P*Δ*DsogMS2. hnt* staining patterns show the opposite, with Δ*PsogMS2* patterns appearing nearly wildtype, and Δ*DsogMS2* embryos showing a pattern similar to, but stronger than, Δ*P*Δ*DsogMS2*. These results suggest that the changes observed in pMAD stainings functionally impacts the subsequent patterning steps, and that changes in the onset and rate of transcription of *sog* have specific and defined consequences in the selection of dorsal fates.

## DISCUSSION

In this study we sought to understand how two shadow enhancers collectively contribute to the output of a gene. We utilized fixed and live imaging techniques to characterize the position, timing, and rate of transcription of each enhancer separately. Far from being redundant, we found these enhancers contributed to different aspects of transcription, and loss of enhancers produced different downstream consequences for development in terms of altered tissue patterning and embryo survivability. Additionally, by separating out different key features of transcription, we have shown that enhancer additivity functions differently at particular steps in transcriptional activation and repression.

### Shadow enhancers show positional preferences along the D/V axis

Our fixed imaging experiments demonstrated that the proximal and distal enhancers contribute to the ventral and dorsal locations of the *sog* transcriptional pattern, respectively, with the highest overlapping activity located in the lateral region of the pattern (Fig. 2). Higher rates of monoallelic expression were seen on both edges of the *sog* pattern in *WTsogMS2* embryos, which are presumably the result of reduction in the frequency of activation the farther away a given nucleus is from the target region of *sog* expression encoded by the enhancers. This is supported by the observation that the peak of monoallelic expression found at either end of the pattern disappears when the enhancer that has a preference for that position is lost. However, it is unclear whether monoallelic expression represents a complete loss of activity from a single allele, or if a small amount of activity remains, but has dipped below our detection threshold for nascent transcription.

### Shadow enhancers interact to mediate repression

Modeling of enhancer activity found that the collective action of the two enhancers complement each other in mostly an additive fashion, that is, the action of the two enhancers together can be adequately explained by assuming that there is no mechanistic interaction between them. However, this is not the case in the ventral portion of the D/V axis where Sna acts to repress transcription of *sog*. Instead, there appears to be enhanced repression by the proximal enhancer in the presence of the distal enhancer, as seen in Fig. 5C where the prediction of our model deviates from the observed *WTsogMS2* data, indicating interaction between the two enhancers.

The cause of this effect is unknown, but a plausible mechanism can be postulated based on the current understanding of Sna-mediated repression. Sna works to repress transcription in the early embryo by the recruitment of the co-repressor dCtBP, which is thought to operate at small genomic distances less than 200bp (Keller et al., 2000). In the classic example of the short range repressive effect of dCtBP, Krüppel is responsible for repressing the activity of the *eve* stripe 2 enhancer to create the sharp posterior border of stripe 2. Located just 1.7kb away is the *eve* stripe 3+7 enhancer, which does not experience any repressive effects despite *eve* stripe 3 being found in the domain where Krüppel is most active in the blastoderm embryo (Nibu et al., 1998). Importantly, the portion of the enhancer that drives stripe 3 is locally depleted for Krüppel binding sites (Vincent et al., 2018).

However, this lack of a shared repressor responsible for recruiting dCtBP is not the case for the enhancers of *sog*, where both enhancers contain binding sites for Sna (see Fig. 1B). Efficient recruitment of the co-repressor by high occupancy of Sna at the distal enhancer may amplify the action of Sna at the proximal enhancer by increasing the local concentration of dCtBP in the microenvironment of the *sog* locus, thereby allowing Sna at the proximal enhancer to recruit dCtBP more efficiently. A modeling based approach that attempted to derive how enhancer sequence changes transcriptional output based on the binding characteristics of recruited transcription factors in *Drosophila* embryos found that Sna repression required uniquely high levels of homotypic cooperativity in the context of a single enhancer compared to all other repressors examined by the study (Fakhouri et al., 2010). It is unknown whether this cooperativity could scale to larger genomic distances, but repressive factors, including the *Ciona* Sna homologue, have been shown to form condensates that may extend the range of repressive activity (Treen et al., 2021).

### Shadow enhancers follow a “first come first serve” model for activation

In the case of activation, our data does not support any mechanism of super-additivity. Activation rates of *sog* are well predicted by a model that assumes enhancers act independently. Decreases in measured t_on_ values seen in *WTsogMS2* embryos are likely accounted for by the wide distribution of t_on_ times measured in Δ*PsogMS2* embryos (Fig. 4C). Activation of *sog* by the distal enhancer occasionally precedes the proximal enhancer, thus modestly lowering the average t_on_ values in *WTsogMS2* embryos. However, in most cases, the proximal enhancer will activate first, and the later t_on_ value contributed by the distal enhancer will be “masked” and will therefore not contribute to raising the average t_on_ value. Because of this, we believe our data supports a “first come, first serve” model of enhancer activation.

Although we do see evidence that RNA Pol II loading rates are diminished in the *WTsogMS2* embryos when compared to Δ*PsogMS2* embryos (Fig. 4D), potentially suggestive of so-called “enhancer interference” (Fukaya, 2021), we believe that this result is well explained by the initial activation of transcription being performed by the proximal enhancer in the majority of nuclei, which appears to drive much lower rates of transcription. This confounds our loading rate measurement, as the rise in signal intensity in *WTsogMS2* embryos is likely a composite of the two enhancers acting sequentially. Techniques that attempt to estimate the promoter state at any given time using an MS2 trace may be able to dissect out the individual contributions each enhancer makes, however we believe that this analysis is not required to explain our data.

### Altered downstream signaling is well predicted by differential transcription activity of shadow enhancer mutants

Our study has uncovered the primary biologically relevant transcriptional parameters responsible for the phenotypic differences in the downstream signaling pathway of *sog*. The slower activating distal enhancer drives insufficient levels of *sog* to achieve the early refinement of the pMAD gradient. However, the high loading rates achieved by the distal enhancer enable enough build-up of Sog in the later stages of NC14 to eventually reach near wildtype restriction of the pMAD domain. In contrast, the faster acting proximal enhancer is capable of achieving an early contraction of the pMAD domain but fails to drive sustained expression of *sog* at high enough levels to continue this contraction. The expansion of the expression domain of the “early” pMAD target gene *ush*, but not the “late” pMAD target gene *hnt* seen in Δ*PsogMS2* embryos, while the opposite is seen in Δ*DsogMS2* embryos, give good indication of the validity of this model.

### Evolutionary considerations for shadow enhancer pairs

With this in mind the question naturally arises: why have two enhancers at all, if it is possible to achieve this result with only one? Based on our previous work on the distal enhancer in reporter constructs, we know that placement of the distal enhancer directly upstream of a promoter is capable of driving fast transcriptional activation at high levels (Yamada et al., 2019). Beyond increasing the robustness of transcription as proposed by previous studies of shadow enhancers (Frankel et al., 2010; Perry et al., 2010; Tsai et al., 2019), we believe our data elaborates on the original hypothesis that shadow enhancers act as a source for evolutionary novelty (Hong et al., 2008). In its original conception, the *de novo* creation of a shadow enhancer allows one of these enhancers to drift, potentially adding new functionality without disturbing the core role of the original transcriptional program.

An alternative view to this interpretation is that selection may favor the creation of enhancers that allow for the tuning of individual transcriptional parameters. In our study, loss of a single enhancer produced defined and unique differences in phenotypic outcomes based on the parameter that enhancer was principally responsible for controlling, either the activation speed in the case of the proximal enhancer, or loading rate in the case of the distal enhancer. By keeping these activities separate, mutations in either enhancer will create smaller, but more precise changes in the downstream patterning events, reducing potential pleiotropy that would be present if *sog* was driven by a single enhancer. Overall, this partitioning of enhancer activity would allow for a more defined exploration of the landscape of potential phenotypes during periods of increased selective pressure.

## MATERIALS AND METHODS

### *Drosophila* lines

All flies were grown on standard fly (*Drosophila melanogaster*) cornmeal-molasses-yeast media. FLy stocks used in this study were: *y[1]w[1118*] (used as wildtype flies) and *y[1] sog[S6]/FM7c, sn[+]* (used as a *sog* null allele; Bloomington Stock Number 2497). *zld*^*-*^ embryos were made using UAS-*zld* shmir lines and the Gal4 driver, MTD as previously described (Sun et al., 2015) Flies of the genotype *y[1] w*; P{His2Av-mRFP1}II*.*2; P{nos-MCP*.*EGFP}2* (Bloomington Stock Number 60340) carried two transgenes, one on chromosome 3, *<u>P{nos-MCP</u>*.*<u>EGFP}2</u>*, which expresses the MS2 coat protein (MCP) fused to EGFP under the control of the *nanos* promoter active in oogenesis, and the other on chromosome 2, *P{His2Av-mRFP1}II*.*2*, which expresses RFP-tagged His2Av in all cells under the control of *His2Av*. Embryos from these and CRISPR engineered flies (see below) were collected on yeasted grape juice agar plates, aged, and either fixed or live imaged (see below).

### Generation of engineered *sog* alleles

All engineered fly lines were created through CRISPR-Cas9 mediated homology directed repair. sog enhancer sequences that were deleted are listed below. Transgenic Cas9 flies were co-injected with pCDF5 plasmids encoding guides targeting relevant genomic targets and pGEM-T vectors containing homology repair templates. All injections were performed by BestGene. pGEMT donor DNA vectors were generated from fragments obtained through genomic PCR for homology arms, and sequences subcloned or PCR amplified from existing plasmids. All 24xMS2 loops containing plasmids utilized the MS2 sequence found in the MS2v5(-TAG) vector (Yamada et al., 2019). The neutral spacer DNA in the primary deletion plasmid was generated using the spacer sequence found in Scholes et al. 2019 (Scholes et al., 2019) and was generated as a IDT (Integrated DNA Technologies) gene block. The 3×3P-RFP sequence (Berghammer et al., 1999; Sheng et al., 1997) for the distal deletion (Δ*D*) plasmid was a generous gift from the Desplan Lab. Plasmids were assembled using a combination of restriction enzyme digest and ligation, and Gibson assembly cloning. Primers used to create donor vectors for each fly line are listed below, along with the guide sequences associated with each injection. Plasmid sequences and maps can be found at https://rushlowlab.bio.nyu.edu/

#### *sog* enhancer sequences

Proximal Enhancer (ChrX: 15,624,486..15,625,257): tgaaaatgcaacaacggcagcgaaccaagaaagaaatagtggaaaaaaaaggaaaaaaaaactgcaactcgggaacataatagtatgcaatatacacatacatttatatgcatatataatatatgagtgtagggagtattgggaggggggtttgcaaacaggaaatgcagctaatcaagcgtgtgagttgcaacaaattgcaattgggtgccgctttatggtccatggtccataccacccaatggtctatatacatgggcaggcatccatttgggtatacccgtatctttttggtaagcggcttacggacgccgatgcgtctgcgcagcgcagtgcaggcagcgagcggaagggaattcccgctttccggattaaaactggacacaataataataaaaaaaaaaaaagaaaacggagtgctatgctgtgccgtcgggaatatcccatgtcccgaaaaccctggcgggattagaggtgcgagcaggtcccgcctcggcaccggctggaattctacctgcgattacggggatttccccgcaccatacagccatatagccatatagccatatagacgacacggcgtatgcgcaatggcattggcaacttatgcaatcgcagcggaggtagaaatgtcgaaagcaacaggcaacagttaatacccctttaactaaagattttgactagttcgaactttaaggatatgcgattgaaagtcgattaaaaactaaacctgataaataactcaaataacctattgaaatattgaaaactc

Distal Enhancer (ChrX: 15,646,594..15,647,337): Atttaatcgaaggactgcaatgggcatatacaacaaattctacgataaaggattcaatattgattgttatatgtttatggcagccaattgatgccgactgacctgtgtgtgtgtgtgtgtgtgtgtggaagctcaggatggacagattcccgggtttcagcggaacaggtaggctggtcgatcggaaattcccaccatacacatgtggctataatgccaacggcatcgaggtgcgaaaacagatgcagcctcataaaaggggcgcagataaggtcgcggttgcgtgggaaaagcccatccgaccaggaccaggacgaagcagtgcggttggcgcatcattgccgccatatctgctattcctacctgcgtggccatggcgatatccttgtgcaaggataaggagcggggatcataaaacgctgtcgcttttgtttatgctgcttatttaaattggcttcttggcgggcgttgcaacctggtgctagtcccaatcccaatcccaattccaatccgtatacccgtatatccaatgcattctacctgtcctgggaatttccgatttggccgcacccatatggccacggatgcgtgagagtgctctccgtgcgattctagatcatcgtgggtattcgcagacaatcgggttattgtgccgcattcgatgttggctctttggttttcggaaactctgaccaggttttcggttttcggtttttgattttgggtttttccggccgcatcgtg

#### Primer and guide sequences

*WTsogMS2*

Plasmid Primers:

5’ Homology Arm Forward

5’-gcctggctgtgtgagtgttgtg 5’ Homology Arm Reverse

5’-cgagatctctgtttatacaaagtcttagc 3’ Homology Arm Forward

5’-tgccgaatcgggtaggacgat 3’ Homology Arm Reverse

5’-accggaacgaatatcgaatatgcaattggc Guide Sequences:

5’-taaacagagatctcgggaag* 5’-aaacagagatctcgggaagt*

Δ*PsogMS2*

Plasmid Primers:

5’ Homology Arm Forward

5’-gcctggctgtgtgagtgttgtg 5’ Homology Arm Reverse

5’-cgagatctctgtttatacaaagtcttagc 3’ Homology Arm Forward

5’-tgccgaatcgggtaggacgat 3’ Homology Arm Reverse

5’-accggaacgaatatcgaatatgcaattggc Guide Sequences:

5’-taaacagagatctcgggaag* 5’-aaacagagatctcgggaagt* 5’-gttgggattctgtttatcaa

5’-tgggcaaatagaaacggcgc

Δ*DsogMS2*

Plasmid Primers:

5’ Homology Arm Forward

5’-gtttttatgtccgtctggcgc 5’ Homology Arm Reverse

5’-gatggctaaaatgaataaaatgagttgcta 3’ Homology Arm Forward

5’-gtcatctggtggcacaggac 3’ Homology Arm Reverse

5’-gaaaggaattccacgtattcgctg Guide Sequences:

5’-taaacagagatctcgggaag* 5’-aaacagagatctcgggaagt*

5’-gcagtccttcgattaaatga 5’-ccaccagatgacgcacgatg

*Guides were used in every experiment to insert the 24x MS2 loops

Proximal+distal deletion(Δ*P*Δ*D*) flies were generated by injecting the Δ*D* guide plasmid and donor plasmid on the background of the proximal deletion (Δ*P)* fly line homozygous for transgenic Cas9. Flies expected to contain 3×3P-RFP cassettes were screened for red fluorescence, all other lines were screened via PCR using primers that spanned the MS2 insertion:

MS2 Screen Fwd

5’-tgacgtttgattagccaccagttggg

MS2 Screen Rev

5’-gccaacctcaacttccaatctccg

### Colorimetric *in situ* hybridization

Embryos were collected and aged to be 1-3 hours old at room temperature and dechorionated in Clorox for two minutes. They were then fixed in 4% formaldehyde (1X PBS) and an equal volume of heptane for 25 minutes while shaking vigorously. Devitellinization was performed by pipetting off the bottom fixative phase and adding 4 mL of methanol and shaking vigorously for 30s. Embryos were rinsed in methanol and transferred to ethanol for storage at − 20°C. Hybridization of fixed embryos used a standard in situ hybridization (ISH) protocol and DIG-labeled *sog* cDNA or lacZ RNA antisense probes; hybridized at 55°C overnight). Visualization of the labeled probe was done using anti-DIG-AP (alkaline phosphatase) antibodies (Roche Biochemicals) followed by histochemical enzymatic staining reagents (Roche Biochemicals). Embryos were mounted on slides with Aqua-Polymount (Polysciences) using 1.5 coverslips (Fisher Scientific), and imaged with Zeiss Axiophot DIC optics and a Zeiss Cam and ZEN2012 software.

### Single-Molecule Fluorescent *in situ* Hybridization (smFISH)

Probe sets for smFISH were generated using the online Stellaris (LGC Biosearch Technologies) probe designer. *sog* probes were ordered to be conjugated to Atto-670, and MS2 probes were ordered to be conjugated to Atto-570. Embryos were fixed in the same manner outlined above, and stained following the *Drosophila* whole embryo staining protocol found on the Stellaris website (https://www.biosearchtech.com/support/resources/stellaris-protocols). After *in situ* staining, embryos were washed 3x with PBS-Tris, and stained overnight at 4 degrees C with anti-Dorsal antibodies (see below) followed by staining with fluorescently labeled secondary antibodies for 1.5 hr at room temperature (see below).

### Antibody staining

Antibody staining was performed at 4 degrees C for 16 hours followed by three 20 minute washes in PBS + 0.1% Tris pH 7.0. Anti-Dl antibody (Dl_7A4) was obtained from the Developmental Studies Hybridoma Bank and used at 1:50 dilution. Anti-pMAD antibodies were obtained from Cell Signaling. Embryos were then stained with secondary antibodies: Alexa Fluor 488 anti-mouse or Alexa Fluor 488 anti-rabbit (ThermoFisher Scientific) for 1.5 hours at room temperature and washed in the same manner. After DAPI (D9542, Sigma-Aldrich) staining for 20 minutes, embryos were mounted on microscope slides using ProLong™ Diamond Antifade Mountant (ThermoFisher Scientific) and Number 1.5 glass coverslips (Fisher Scientific). Embryos were imaged with Zeiss 880 with Airyscan confocal microscope.

### Fixed tissue confocal imaging

All confocal images were captured on an LSM Zeiss 880 microscope. Images for the pMAD experiments were captured using a 20X objective with 1.1 Digital Zoom and a 2000×800 scanning area. Images all contained approximately 20 Z-planes. Laser power was set for lasers 405nm at 0.5% and 488nm at 3%, with gain set at 750. Images for all smFISH experiments were captured using the Airyscan module, and processed using the suggested Airyscan Processing strength. These images were captured using a 40X objective with 1.0 Digital Zoom and a 2000×1500 scanning area. Images all contained approximately 50 Z-planes. Laser power was set at 0.5% (405nm), 5% (488nm), 7% (561nm) and 20% (633nm), with gain set at 750.

### Live confocal imaging

Virgin females maternally expressing MCP-GFP and H2Av-RFP were crossed with males of the MS2 reporter lines. 0-1 hour embryos were collected, dechorionated, and transferred onto a breathable membrane (Lumox Film, Sarstedt AG & Co.) in the middle of a plastic microscope slide (3D printed on Ender 3 Pro, Creality). Live imaging was performed using a LSM Zeiss 880 63X objective lens with the following settings: optical sections: 1024×1024 pixels, 20z stacks 0.7μm apart, 12bit; zoom: 1.0; time resolution: 25 seconds per frame. Laser power was set at 0.6% (488nm), and 0.4% (561nm) with gain set at 800. Embryos were imaged for approximately one hour, typically from NC12 to late NC14.

### Image analysis, quantification and statistical analysis

Processing for images followed a pipeline started with feature extraction using standard tools in Imaris, then data exported to .csv files for organization, further processing, and plotting. For pMAD experiments, nuclei positions were obtained using the “spots” function with an estimated diameter of 4um, and a Z-axis diameter of 7um, with background subtraction enabled. Spot positions were restricted to an area of interest approximately 75% to 25% of egg length. Fluorescence intensity from the pMAD channel at all spot positions was extracted and processed using the “pMAD_quant.R” script. This script aligned, plotted, and extracted gradient widths from all pMAD gradients measured.

For smFISH experiments, nuclei positions were instead obtained using the “volume”, with a surface detail parameter set at 0.2um, and background subtraction enabled. Foci of *sog* smFISH signal were obtained using the “spots” function, with an estimated diameter of 0.5um and a Z-axis diameter of 1. Alleles were discriminated by analysis of MS2 signal at spot locations, and thresholds were set manually by examining the separation between the two populations. Foci were assigned to single nuclei by finding the nearest nucleus in 3D space to each focus. Nuclei with more than two assigned foci were excluded from the analysis, and represented less than 1% of the data.

Live imaging analysis was performed on Imaris by tracking nuclei using the “spots” function with an estimated diameter of 4um and a Z-axis diameter of 6um. Tracking was performed using the “retrograde motion,” with a max allowable gap of 1, and a max allowable displacement of 10um. Foci were also tracked using the “spots” function with an estimated diameter of 1.3um and a Z-axis diameter of 2um. Tracking was performed using the “retrograde motion,” with a max allowable gap of 0, and a max allowable displacement of 2.5um. Spots were filtered by inclusion of foci with “Quality” scores greater than 33.0, median RFP fluorescence greater than 200 AU, mean GFP fluorescence greater than 250 AU, and a distance from the xy-border greater than 1um. All tracking data, including position and mean GFP fluorescence was exported to .csv files for further analysis in R.

Foci were assigned to nuclei by finding minimum distance between foci and nuclei. Subsequently, any nuclei that came within 3um of the xy-border were filtered out to reduce edge effects. Nuclear cycle times and D/V axis relative positions to the mesoderm were annotated manually and stored in a separate .csv file. Nuclei were assigned into positional bins by taking the difference between the annotated mesoderm y-coordinate and the average position of the nucleus for each nuclear cycle. t_on_ values for NC13 and NC14 were obtained by subtracting the time GFP foci were first detected from the annotated cycle time of the respective nuclear cycle. Loading rates were estimated by fitting a linear model to the first five timepoints of the GFP foci intensity. Negative values were discarded, and represented less than 5% of the data. Total output values were calculated by multiplying each nucleus’ loading rate by the total time that foci was detected, with a max allowable time of 25 minutes in NC14 to account for differences in imaging time between each movie. These values were then averaged for each positional bin, and multiplied by the percentage of active nuclei in the corresponding bin.

### Plotting

All plots were generated using base R plotting functions. All error bars were computed using the standard error of the mean (s.e.m.).

### Modeling

Models of activation were constructed by fitting gamma distributions to measured t_on_ and t_off_ values using the function fitdist() included in the “fitdistrplus” library. Fits were achieved via maximum likelihood estimation. The shape and rate of each distribution was extracted, and used to construct new distributions of values which were sampled independently to generate simulated t_on_ and t_off_ values. Distribution parameters were subsequently refined by comparing simulated nuclei to measured activation traces for each bin. New sets of potential shapes and rates for each distribution were generated by allowing each parameter to vary by up to 20%, and selecting new shape and rate values based on which parameters minimized the residuals between the prediction generated by the model and data.

During simulation, each nucleus was assigned a t_on_ value and t_off_ value generated from the corresponding distribution. At each timepoint, the number of nuclei that had a t_on_ value less than the current time, and a t_off_ value greater than the current time were considered “on”. The number of nuclei “on’’ was divided by the total number of nuclei in the simulation, generating the value of the proportion of nuclei active for that timepoint. If the assigned t_off_ value was less than the assigned t_on_ value, the nucleus was considered “off” at every timepoint. This allowed us to account for nuclei which never activate transcription without skewing the distribution of t_on_, which was critical to accurately simulate the ventral bins.

For the combined model, nuclei were assigned two t_on_ and t_off_ values each sampled from the two different enhancer deletion distributions. Nuclei were evaluated in the same manner as described above, but only required one enhancer’s values to meet the criteria of “on” to be considered as such. All simulations were carried out using a set of 10,000 nuclei, which represented a compromise between accuracy of prediction and computing power.

## Acknowledgements

The authors would like to thank Sevinc Ercan for many insightful suggestions over the course of this study, and Enrique Rojas for helpful discussions of our model. The authors also thank Michelle Lanis Pollack, Alicia Lina Zhu, Yandel Morel, Lauren Stafford, and Dhruvansh Shah for their tireless help with embryo collection and PCR screening for crispants, and Caichen Duan for help with the live imaging.

## Author Contributions

All authors contributed to the experiments. PHW and CAR designed the study. PHW carried out the image analysis and modeling, and prepared manuscript figures and first draft. PHW and CAR revised the manuscript.

## Data and Code Availability

Imaris generated .csv files and R scripts can be found at: https://rushlowlab.bio.nyu.edu/research/

For any detailed procedures for required file headers or help implementing these scripts please contact the Rushlow lab directly.

